# Inactivation of SARS-CoV-2 by β-propiolactone Causes Aggregation of Viral Particles and Loss of Antigenic Potential

**DOI:** 10.1101/2021.04.22.441045

**Authors:** Divya Gupta, Haripriya Parthasarathy, Vishal Sah, Dixit Tandel, Dhiviya Vedagiri, Shashikala Reddy, Krishnan H Harshan

## Abstract

Inactivated viral preparations are important resources in vaccine and antisera industry. Of the many vaccines that are being developed against COVID-19, inactivated whole-virus vaccines are also considered effective. β-propiolactone (BPL) is a widely used chemical inactivator of several viruses. Here, we analyze various concentrations of BPL to effectively inactivate SARS-CoV-2 and their effects on the biochemical properties of the virion particles. BPL at 1:2000 (v/v) concentrations effectively inactivated SARS-CoV-2. However, higher BPL concentrations resulted in the loss of both protein content as well as the antigenic integrity of the structural proteins. Higher concentrations also caused substantial aggregation of the virion particles possibly causing undesirable outcomes including a potential immune escape by infectious virions, and a loss in antigenic potential. We also identify that the viral RNA content in the culture supernatants can be a direct indicator of their antigenic content. Our findings may have important implications in the vaccine and antisera industry during COVID-19 pandemic.

## INTRODUCTION

The ongoing COVID-19 pandemic caused by the coronavirus SARS-CoV-2 is devastating the human lives across the globe [1, 2]. As of the middle of April 2021, the virus is estimated to have infected about 140 million people, killing over 3 million of them worldwide. The disease is characterized by acute respiratory illness resulting in severe breathing difficulties very similar to the common flu accompanied by severe cough in the infected people forcing several of them to be hospitalized. In addition, there have been several reports of sepsis, blood clotting and multi-organ failure in several persons infected by the virus. The severity of the disease is significantly higher in persons with co-morbid conditions such as diabetes, hypertension, cancer and respiratory problems [2, 3]. Currently, there are no drugs that can cure COVID-19. Vaccines are the best hopes for ending this pandemic. Antibody-based interventional therapies are of great importance in treating severe cases. Some of these projects utilize inactivated virus particles that are used for immunization along with adjuvants. Large-scale viral cultures and antigens are essential for successful generation of vaccines and antisera that are based on whole virus-derived antigens. This requires complete inactivation of the virus particles while causing minimum damage to their structural and antigenic properties. Therefore, these projects are dependent on the availability of viral samples that are totally inactivated while retaining their antigenicity sufficient to induce antibody response.

Several methods have been adapted historically to inactivate the viral stocks that are used for vaccine and antisera development. They include physical methods such as heat [4] and γ-rays [5] or chemical agents such as formaldehyde and β-propiolactone (BPL) [6–8]. SARS-CoV-2 is reported to be inactivated by temperatures starting from 56 °C [9, 10]. BPL mediated inactivation of SARS-CoV and SARS-CoV-2 has been demonstrated [7, 9, 11, 12]. Heating is known to denature the proteins that might negatively impact their antigenicity [4]. γ -irradiation has been employed in several vaccine studies, but they are also known to damage the antigens if not properly optimized [13]. Formaldehyde, despite being one of the oldest and most easily available chemical agents to inactivate viruses, also causes loss in protein antigenicity and hence is less desirable. BPL has emerged as a very popular chemical agent in various vaccine initiatives due to its high inactivation potency and relatively low damage of antigens.

BPL has also been used in SARS-CoV-2 inactivation at various concentrations. However, a comprehensive analysis of its potency and impact on the integrity of viral antigen is not available. In this study, we attempt to optimize the concentration of BPL for SARS-CoV-2 for efficient inactivation and protection of antigenicity. We demonstrate that BPL at 1:2000 concentrations (v/v) is enough to inactivate SARS-CoV-2 and increasing the BPL concentration above 1:1000 leads to significant drop in the antigenic potential of viral proteins, probably caused by the modifications on their amino acids. Our investigation also identified a lack of correlation between viral RNA titer and infectious viral units in the supernatant. However, the viral RNA titer showed strong correlation with the antigenic content in the sample. Our studies also demonstrate that SARS-CoV-2 particles tend to form larger aggregates with increasing concentrations of BPL above 1:1000 which could possibly lead to reduced epitope exposure thereby deleteriously affecting the antigenic potential of the sample.

## MATERIALS AND METHODS

### Cell culture and reagents

Vero cells were cultured in DMEM with 10% FBS (Hyclone, SH30084.03) and penicillin-streptomycin cocktail (Gibco, 15140-122) at 37°C and 5% CO_2_. Anti-Spike antibody was procured from Novus Biologicals (NB100-56578) while anti-Nucleocapsid was procured from Thermo Fisher (MA5-29982). HRP-conjugated anti-rabbit secondary antibody was purchased from Jackson ImmunoResearch (111-035-003). BPL was procured from Himedia Laboratories (TC223-100). The zinc staining kit was from G Biosciences (Reversible Zinc Stain; 786-32ZN).

### Virus culturing

The oro- and nasopharyngeal patient swabs transported in VTM were screened using SARS-CoV-2 specific primers (LabGenomics; Labgun COVID-19 RT-PCR kit; CV9032B) and the samples with low Ct (less than 20) values were chosen to culture virus. VTMs were filter-sterilized and added to Vero monolayers in 96-well plate. Three hours post-infection, the media was replaced with fresh serum sufficient media. The infected cells were further incubated at 37°C with 5% CO_2_ in a humidified chamber and cytopathic effects (CPE) were examined every 24 hours. Cells along with the supernatants were collected from those wells displaying CPE and transferred to fresh 12-well plate containing Vero monolayers for further propagation. This process was repeated until the cell culture supernatant showed a Ct value lesser than 20. The supernatants were titrated for infectious particle count by plaque-forming assay. In the case of dry-swab sample, the swab was first soaked in TE buffer for 30 minutes [14] and further stored at −80°C freezers. Later the sample was used as inoculum for infection similar to VTM.

### Virus quantification, titration, and sequencing

RNA from VTMs or swab-immersed TE buffer or cell culture supernatant was isolated using viral RNA isolation kit (MACHEREY-NAGEL GmbH & Co. KG; 740956.250). The SARS-CoV2 RNA was quantified using (LabGun^™^ COVID-19 RT-PCR Kit) following the manufacturer’s protocol or following WHO guidelines using SuperScript^™^ III Platinum^™^ One-Step qRT-PCR Kit (ThermoFisher) and Taqman probes against CoV-2 E, and RdRP (Eurofins Scientific). The isolates that were established in cultures were sequenced by next-generation sequencing and compared with Wuhan SARS-CoV-2 genome as reference S. The sequence of isolates used for all the experiments here were submitted to GISAID public database (GISAID ID: EPI_ISL_458075; virus ID-hCoV-19/India/TG-CCMB-O2-P1/2020, and EPI_ISL_458046; virus ID-hCoV-19/India/TG-CCMB-L1021/2020). Subsequently, the CCMB_O2 isolate was scaled up and used for the experiments. All the virus cultures were titrated for infectious particle count using plaque forming assay (PFU/mL) before use. Briefly, the supernatant was log-diluted from 10^-1^ to 10^-7^ in serum-free media and was added to a 100% confluent monolayer of Vero cells. Two hours post-infection the infection inoculum was replaced with agar media (one part of 1% LMA mixed with one part of 2 × DMEM with 5% FBS and 1% Pen-Strep). 6-7 days post-infection cells were fixed with 4% formaldehyde in 1× PBS and stained with 0.1% crystal violet. The dilution which had 5-20 plaque was used for calculating PFU/mL.

### Virus infection and inactivation

Cells were infected with SARS-CoV2 at 90% confluency in serum-free media at 1MOI for two hours and subsequently, the inoculum was replaced with fresh serum-free media. Three days post-infection, cell culture supernatant was collected and the debris was removed by centrifugation and stored until further use. Later, the infectious viruses in the supernatants were inactivated using BPL at varying concentrations (1:250, 500, 1000, 2000 (v/v to the culture-media). In brief, the supernatant with BPL was incubated at 4°C for 16 hours followed by 4-hour incubation at 37°C to hydrolyze the remaining BPL. The inactivation of the virus was confirmed by the absence of CPE in three consecutive rounds of infections. To study the effect of BPL on viral antigenicity, either live infectious or inactivated viral samples were concentrated to 10 × using centrifugal filter units with 100 kDa cut-off.

### Zinc Staining

All the reagents provided by the manufacturer (Reversible Zinc Stain; G Biosciences) were diluted to working concentration using deionized water. Zinc staining was done after performing SDS-PAGE under reducing conditions. Viral supernatants concentrated either by ultracentrifugation or by centrifugal filters were lysed with equal volume of 2 I × lysis buffer (containing 2% NP40, 100 mM Tris-HCl, 300 mM NaCl, 2 mM sodium orthovanadate, 2 mM phenylmethylsulphonyl fluoride, 20 mM sodium pyrophosphate, and protease inhibitor cocktail) [15], mixed with 6 × Laemmli buffer, boiled and loaded onto the gels after cooling. Gels were washed with distilled water after electrophoresis followed by incubation in 25 mL washing buffer (Reagent I) for 5 minutes on a shaking platform. Subsequently, the gel was incubated with 25 mL of Reagent II containing imidazole for 15 minutes. Finally, the gel was stained in zinc sulfate-containing Reagent III for 45-60 seconds followed by immediate transfer to distilled water. The gel image was scanned against a dark background and was then destained using destaining solution provided by the manufacturer followed by three 5-minute washes using distilled water. The destained gel was used for subsequent western blot transfer procedures.

### Immunoblotting

Fresh gels or the destained gels after zinc staining were transferred onto PVDF membranes for 16 hours at 30 V. Membranes were blocked in 5% BSA and were subsequently probed with SARS-CoV-2-specific Nucleocapsid (1:8000) or SARS Spike (1:2000) antibodies. HRP-conjugated goat anti-rabbit secondary antibody was used at 1:20000 dilutions. Blots were then developed with ECL reagents (Clarity ECL Western Blotting; Bio-Rad) using ChemiDoc MP system (Bio-Rad). All densitometric analyses were performed using ImageJ software [16].

### Dynamic light scattering spectroscopy

The particle size of virions from BPL inactivated viral samples was measured using a dynamic light scattering instrument (SpectroSize 300; Nabitec). The instrument uses a laser diode and operates at a wavelength of 660 nm at 90° scattering angle and detection was done by an avalanche photodiode detector. Each BPL inactivated sample was subjected to twenty measurements to obtain the particle size. The data obtained was analyzed using SpectroSize 300 software that provided the average hydrodynamic size distribution profiles. The average size of the particles from each sample was plotted against the dilution of BPL used. The experiment was repeated in triplicate using samples with different Ct values.

### Statistical analysis

Viral supernatants used in DLS were independently inhibited with BPL and were considered as biological replicates. At least three independent replicates were used. c to generate mean ± SEM, which are plotted graphically. To calculate statistical significance, two-tailed unpaired student *t*-test was performed and the resultant *p* values were represented as *, **, *** indicating *p* values ≤ 0.05, 0.005, and 0.0005 respectively.

## RESULTS

### Establishment of SARS-CoV-2 cultures

Several other articles have described methods for the establishment of SARS-CoV-2 cultures [17–20]. Here, the key is to have access to patient oro- or nasopharyngeal samples in viral transport medium (VTM) that display low Ct values in the quantitative real time RT-PCR assays. The association of the viral load estimated by qRT-PCR with COVID severity is controversial [21, 22], and qRT-PCR-generated Ct values of viral genes are not true indicators of the viral load owing to a large number of variables in sample collection and processing and hence could be deceptive at times. Therefore it is important to try several VTM samples that display Ct values below 30. Even though lower Ct values are indicators of high viral load, several samples with low Ct values could not establish a culture. This could primarily be determined by the infectious viral load in the sample that is dependent on the collection process and the post-collection storage conditions. Samples that did show signs of infection in a small scale (96-well set up) were then gradually expanded to larger scale to maintain a constant viral culture for further experiments. Each passage of virus was tested for viral load and titer to ensure the retention of infectivity (Table 1).

**Table 1:**
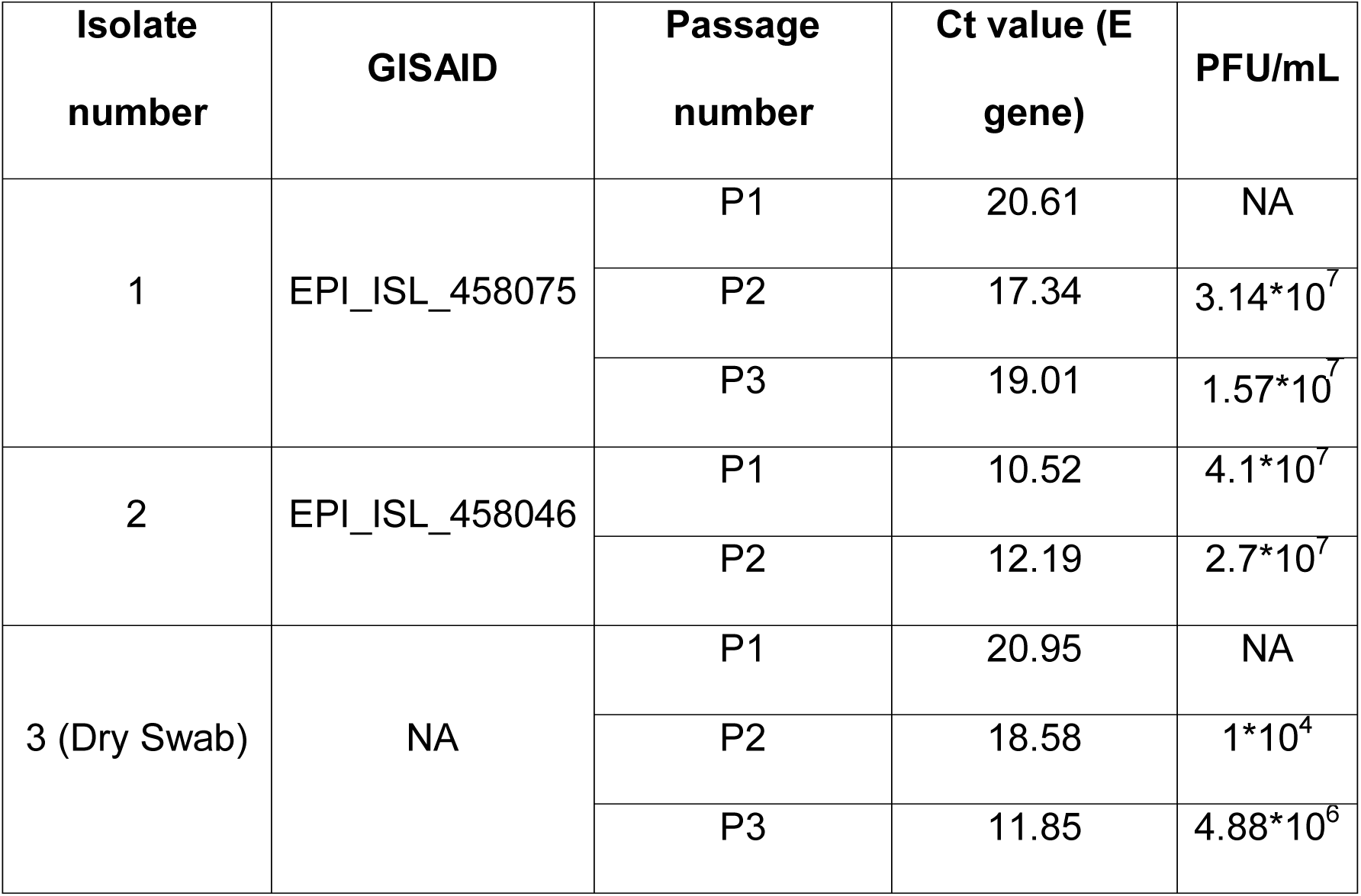
Viral RNA contents and titers of three independent SARS-CoV-2 preparations isolated from patient swab samples and passage to larger formats. “P” indicates the passage number. The culture in which Vero cells were incubated with the patient sample was designated as P1. The third sample was isolated by the dry swab method as mentioned.

Recently, RNA extraction from the swab samples was shown to be dispensable thereby enabling direct processing of the samples for qRT-PCR in COVID-19 screening [14] as well as in other respiratory viral infections [23, 24]. Collecting the samples in dry-swab form has been demonstrated to be effective in preserving the viral content and also much more bio-safe [23, 24]. The dry swabs were immersed in TE buffer before using directly in qRT-PCR. However, the dry-swab method has a potential risk of inactivating the virus due to long-term dry conditions. We tested if virus cultures can be established from the dry-swab samples resuspended in TE buffer stored at −80°C. As in the case of VTM, the potential samples were incubated with Vero cells for infection. Interestingly, as demonstrated in Table 1, we successfully isolated SARS-CoV-2 from dry-swab collection sample, indicating that virus particles derived from dry-swab method can be viable and can indeed establish infection.

### Ct values do not correlate with infectivity, but with protein content

Several studies have used Ct values as a measure of viral titer in the culture supernatants [25, 26]. During our studies, we encountered numerous instances where low Ct values do not really translate into a high viral titer (Figure 1A). Supernatants with Ct values differing significantly, displayed comparable viral titers. We often came across samples with low Ct values and low infectious titers and also those with relatively higher Ct but with high viral titers. Large majority of the established culture supernatants had infectious titers around 10^7^ PFU/mL but their Ct values ranged from 10-28, clearly pointing to a substantially large fraction of viral RNA contributed by non-infectious viral particles or non-virion associated viral RNA. However, samples with low Ct values corresponded to high viral protein content (Figure 1B) suggesting that the samples with low Ct values and low titers contained larger amounts of defective and noninfectious viral particles that contributed to the higher protein content. Thus, the samples with low Ct values are ideal for the preparation of antigens irrespective of their infectious viral units.

**Figure 1.**
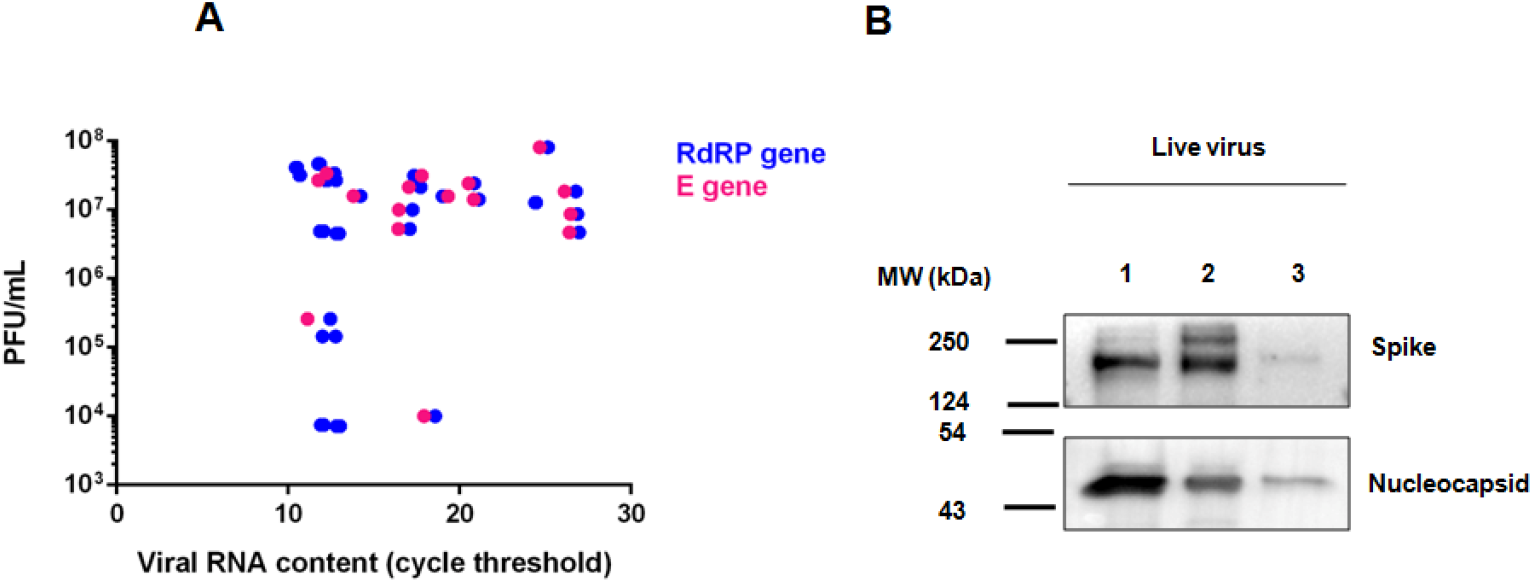
RNA content of the virus cultures need not be correlated with the infectious titers, but with the antigenic content. (A) Viral RNA content in cultures by qRT-PCR of E and RdRP genes were plotted against the infectious titers of the same cultures. RNA samples prepared from the supernatants were subjected to qRT-PCR for the detection of SARS-CoV-2 E and RdRP genes. Infectious titers of the cultures were determined by plaque forming assays. (B) Analysis of the relative protein content of three viral supernatants with different viral RNA contents by immunoblotting of viral proteins. Sample 1, 2 and 3 had Ct values of 12.8, 14 and 17.7, respectively as determined by qRT-PCR of E gene. Immunoblotting was performed using specific antibodies.

### Inactivation of SARS-CoV-2 by BPL

BPL is a common reagent used for chemical inactivation of viruses [6, 27, 28]. We titrated the optimal concentration of BPL required for inactivation of SARS-CoV-2. BPL was added to viral supernatants with known infectious titer (PFU/mL) to make final dilutions of 1:2000, 1:1000, 1:500 and 1:250 (all v/v). Infectivity of the supernatants was measured by CPE. Our results demonstrate that BPL was consistently effective in inactivating the virus completely even at 1:2000 dilutions. Three consecutive rounds of infection confirmed total inactivation of virus at these concentrations (Table 2; Figure 2). These results indicate that BPL at 1:2000 concentrations is strong enough to inactivate SARS-CoV-2 efficiently.

**Table 2.** Demonstration of complete inactivation of virus particles by BPL after three rounds of consecutive culturing.

| Sample ID | Before Inactivation |  | Inactivated virus infectious titer after three rounds of consecutive culturing |  |  |  |
| --- | --- | --- | --- | --- | --- | --- |
|  | Ct (E gene) | Infectious titer (PFU/mL) | 1:250 | 1:500 | 1:1000 | 1:2000 |
| Sample 1 | 12.79 | $2.71 \times 10^7$ | - | - | - | - |
| Sample 2 | 17.72 | $2.14 \times 10^7$ | - | - | - | - |
| Sample 3 | 25.16 | $2.71 \times 10^7$ | - | - | - | - |

**Figure 2.**
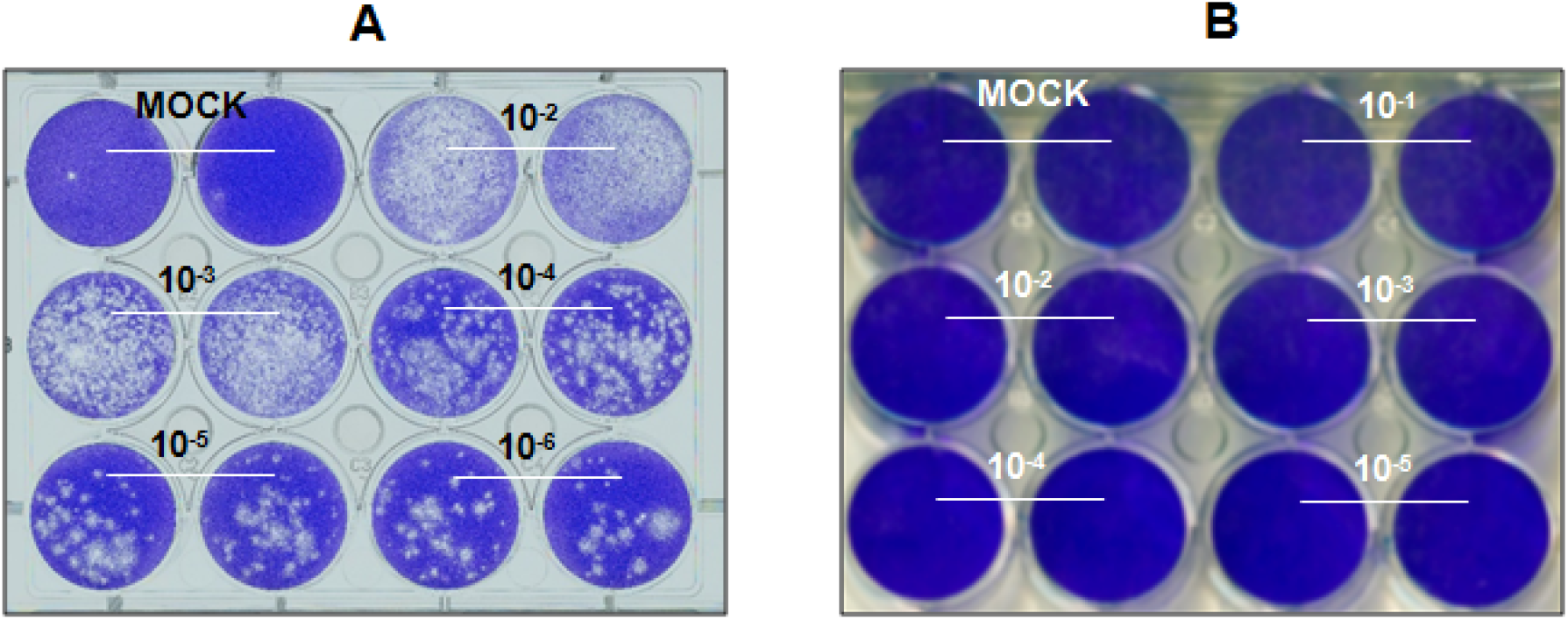
Confirmation of the presence of infectious SARS-CoV-2 particles in the supernatant of the cultures. (A) Representative PFU assay plate showing consistent drop in the plaques with logarithmic dilution of the sample. (B) Confirmation of the inhibition of infectious virus particles with BPL treatment. Viral supernatant treated with BPL at 1:250 concentrations were inoculated with Vero cells and CPE was monitored for six days. Untreated supernatants were used in the control experiments. After six days, the supernatants were further inoculated with fresh cells and this was repeated for a third round of infection. The image is from the third round of repeated infection demonstrating the absence of CPE, confirming total inactivation of infectious virus particles.

### BPL inactivation causes damage to SARS-CoV-2 antigens

One of the major requirements of BPL inactivation is in vaccine studies. However, BPL treatment consistently interfered with the quantification of viral protein. BPL is known to damage the genetic content, but its influence on the proteins of the virions is less understood [29]. In addition, if BPL damages structural proteins of the virions, this could be less desirable for vaccine studies. To address this issue, we tested the epitope integrity of SARS-CoV-2 virions inactivated with BPL. Viral supernatants treated with BPL at 1:250 dilutions were concentrated 10 × by filters with 100kDa cut-off membranes. Protein lysates prepared from these concentrates were electrophoresed by SDS-PAGE following which the gels were zinc stained to visualize antigens. BPL treatment caused minimal drop in the total protein content in the samples as demonstrated in Figure 3 A and B. Next, we studied the effect on antigenicity by detecting structural proteins spike (S) and nucleocapsid (N) by immunoblotting. Immunoblotting against S and N was used as a proxy measure of the antigenic integrity of the virions. Any drop in the band intensity would be considered as the outcome of potential damage to the epitopes. The immunoblots revealed a substantial loss of signals in BPL treated samples against the untreated, infectious samples (Figure 3 C and D) suggested that BPL treatment is causing the loss of antigenic integrity in addition to causing loss in the protein content.

**Figure 3.**
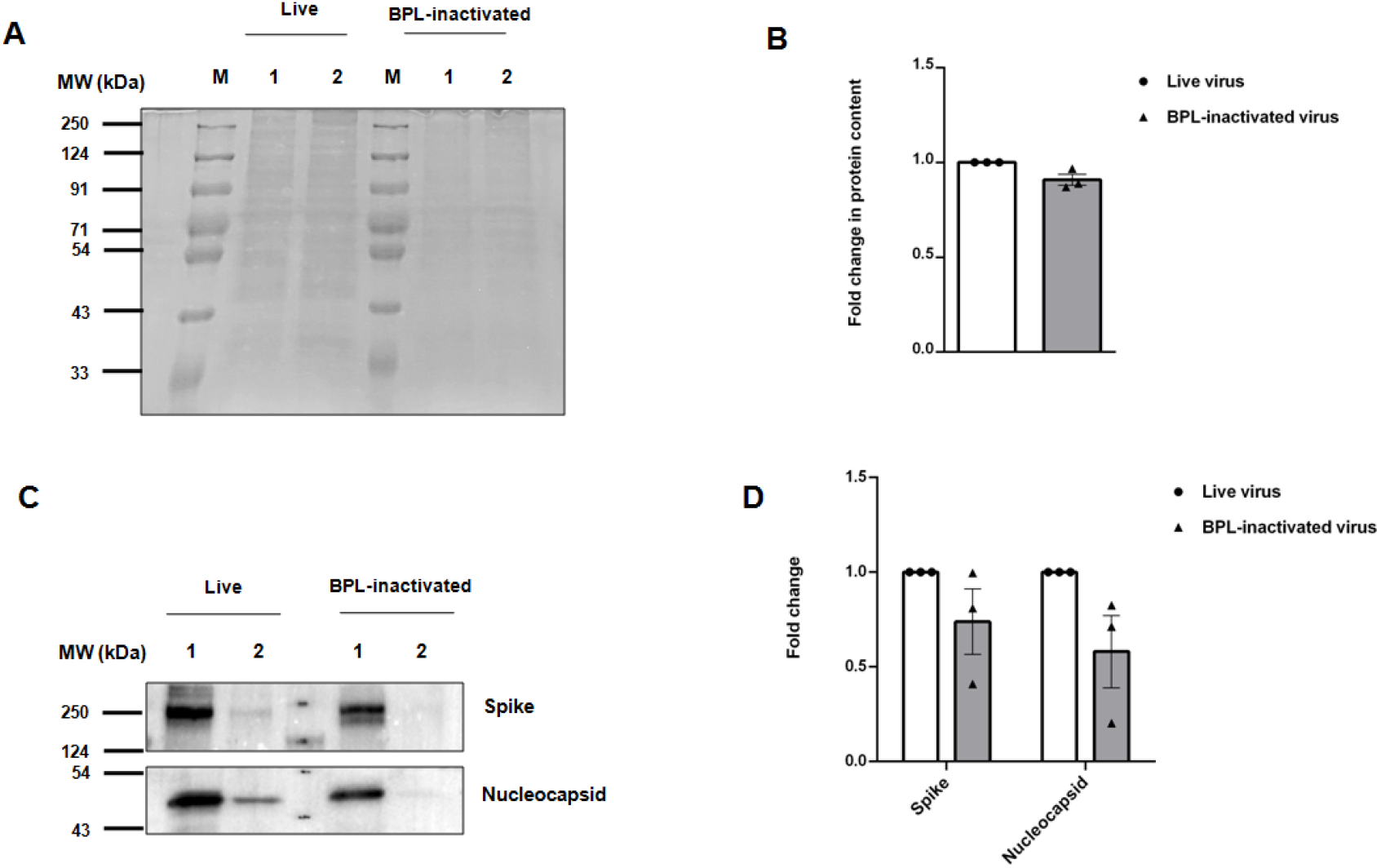
Qualitative analysis of the BPL inactivated virus samples with the infectious control samples. (A) Zinc stain of SDS-PAGE with samples from two individual live infectious or their corresponding inactivated samples. M indicates protein molecular weight marker and the numbers indicate two individual samples. Inactivation was performed by BPL at 1:250 concentrations. The concentrated samples were lysed in equal volumes of 2 × protein lysis buffer, mixed with 6 × Laemmli buffer, boiled and loaded into the gels. After electrophoresis, the gels were stained following the instructions provided by the manufacturer. (B) Relative protein contents in the active and inactivated samples by ImageJ analysis. (C) Immunoblots for SARS-CoV-2 Spike and Nucleocapsid proteins. Concentrated samples separated on SDS-PAGE were transferred onto PVDF membrane and subsequently immunoblotted using specific antibodies against the proteins. (D) Relative antigenic integrity in infectious and BPL inactivated virus samples. Immunoblot images were analyzed by ImageJ software for quantification.

In order to further substantiate this point, we treated the viral supernatants with varying concentrations of BPL at 1:2000, 1:1000, 1:500 and 1:250 dilutions. Supporting our initial observations, loss in the signal was the most striking in sample with 1:250 BPL concentrations followed by 1:500 (Figure 4A). BPL caused much less loss at 1:1000 and 1:2000 dilutions. This was further strengthened by immunoblotting where a gradual loss of antigenic integrity was remarkably captured in 1:250 dilutions (Figure 4B). This could be caused either by a possible chemical modification of amino acids in the epitopes or by the potential loss of exposure of epitopes caused by aggregation. These results collectively demonstrate that BPL treatment causes loss in protein content and also a further loss in antigenicity and suggest that using 1:2000 or 1:1000 dilutions would be more appropriate for vaccine studies.

**Figure 4.**
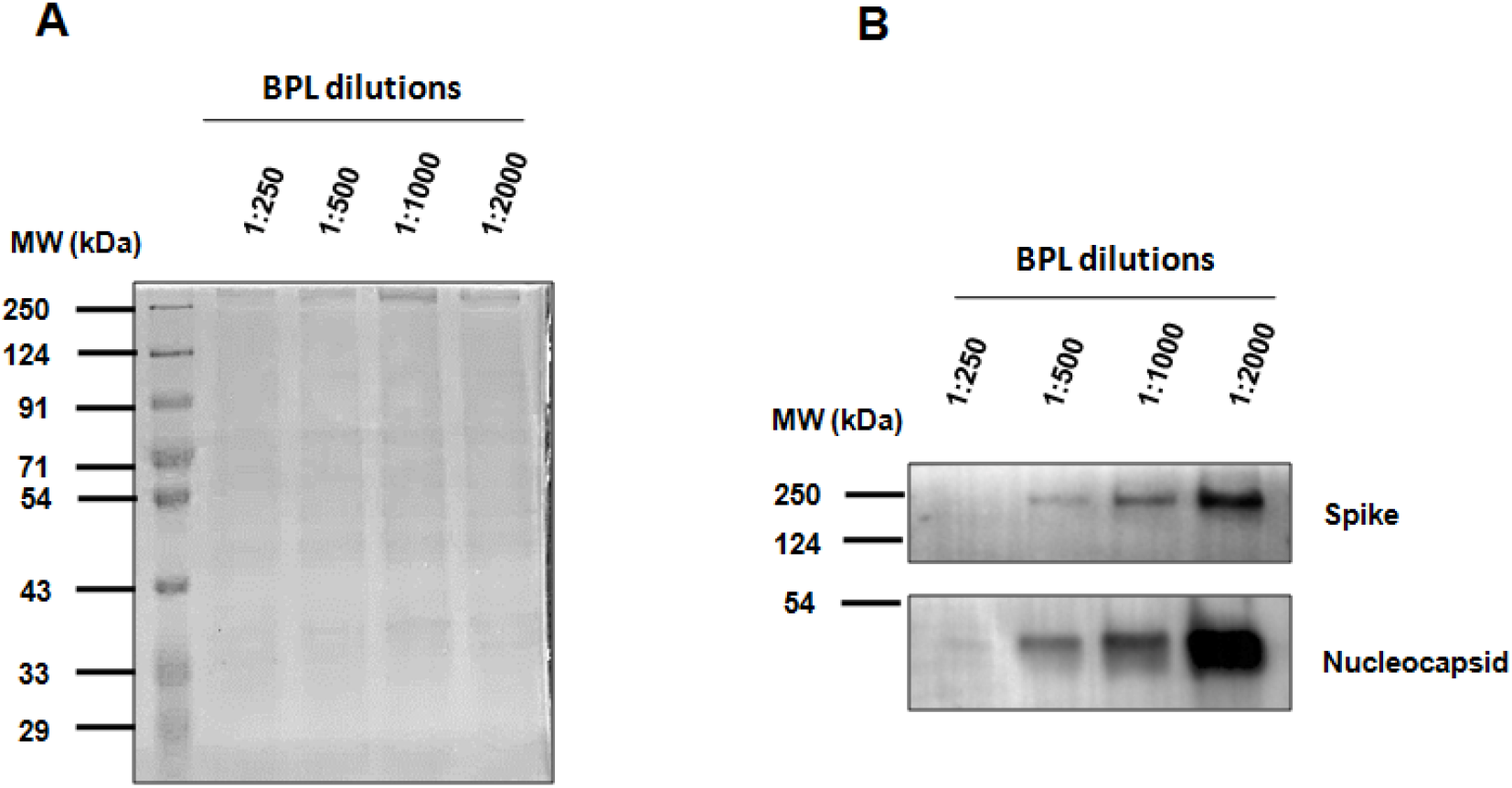
Confirmation of the loss of antigenic integrity by BPL (A) Viral supernatants were treated with varying concentrations of BPL as mentioned in the figure and subsequently concentrated before separating on SDS-PAGE. As in Figure 3A, the gel was zinc stained to visualize the protein bands. (B) Immunoblotting of the viral proteins in samples treated with varying concentrations of BPL. Samples were separated on SDS-PAGE after which they were transferred onto PVDF membrane and immunoblotted.

### BPL treatment causes aggregation of SARS-CoV-2

Earlier studies have demonstrated that BPL treatment causes aggregation of virus particles [6]. To test whether SARS-CoV-2 undergoes aggregation during BPL treatment, we used dynamic light scattering (DLS) that can study particle size distribution in a suspension. Viral supernatants treated with varying concentrations of BPL were analyzed by DLS. Interestingly, increasing concentrations of BPL induced the formation of larger aggregates as demonstrated in Figure 5 A-D. While the viral particles in the supernatant with BPL at 1:2000 dilutions had an average size of about 160nm their size gradually increased to over 500nm with 1:250 concentration of BPL (Figure 5E). These results demonstrate that BPL causes aggregation of SARS-CoV-2 particles in a concentration-dependent manner. Increased aggregation of virions could result in significant loss in the exposure of the epitopes and hence would render them less suitable for antibody response. Additionally, filtration of the mixture post-BPL treatment is not advisable as it might cause significant loss of virion aggregates. Increased aggregation coupled with lower exposure of viral proteins indicates that higher concentrations of BPL are not optimal for inactivation of virus.

**Figure 5.**
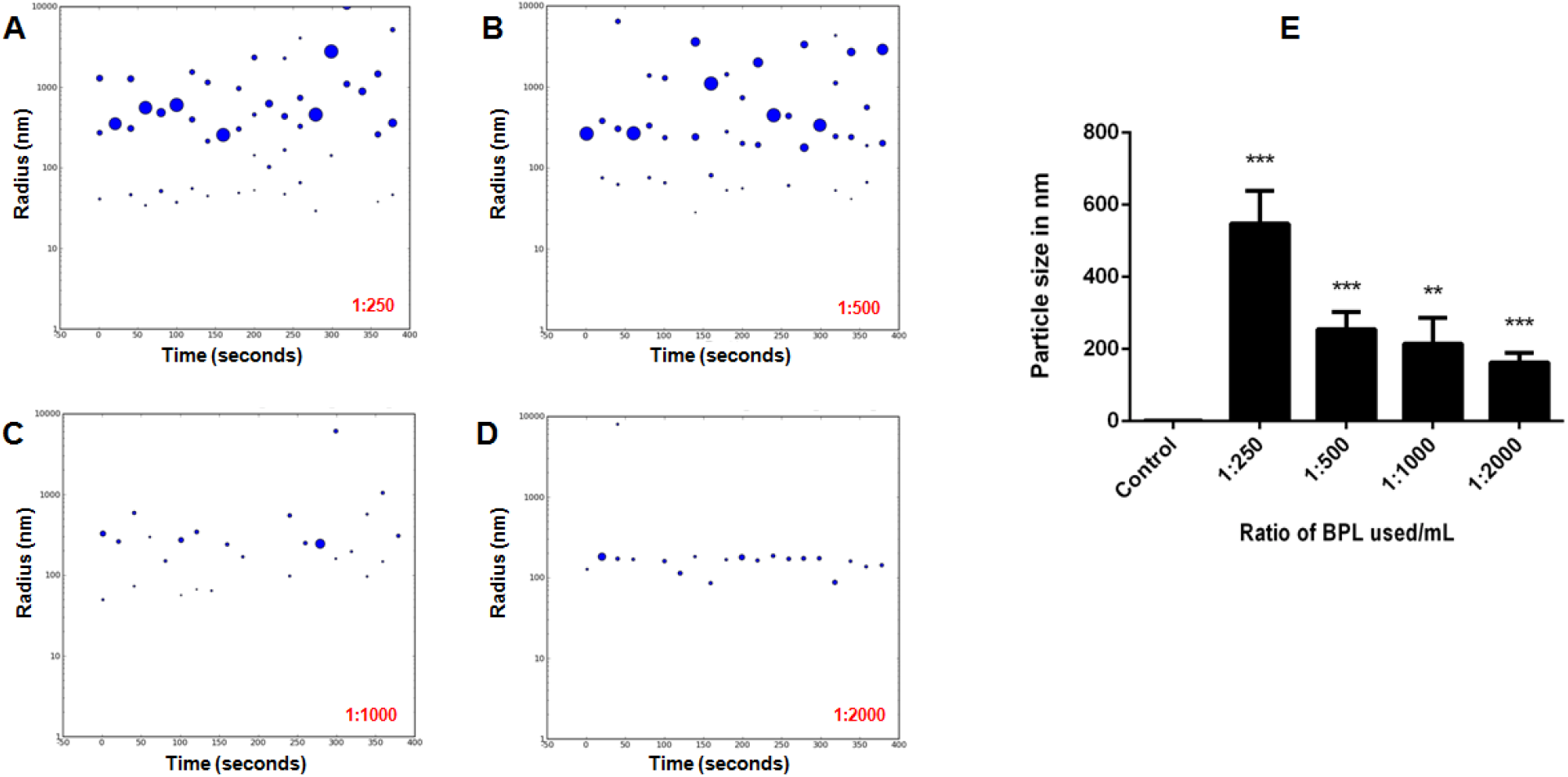
BPL causes aggregation of SARS-CoV-2 particles. (A-D) Representative images of analysis of the particle size of SARS-CoV-2 inactivated with BPL at 1: 250, 1:500, 1:1000 and 1: 2000 concentrations (v/v) by dynamic light scattering. (E) Average size of the virus particles treated with varying concentrations of BPL as mentioned.

## DISCUSSION

In this study, we focused on characterizing the methods for preparation of large volumes of inactivated SARS-CoV-2 cultures for therapeutic purposes such as vaccine and antisera production. Since BPL is the mode of choice for inactivation of several microbes, we optimized the concentration and studied the impact of the treatment on the epitopes and virus aggregation. We demonstrate that BPL at 1:2000 (v/v) dilutions in the culture supernatant is sufficient to totally inactivate the virus. Our studies suggest that BPL negatively impacts the antigenic potential of the virus thereby potentially affecting the immune response when used as antigens. However, lower concentration of BPL at 1:2000 concentrations had minimal impact on the antigenic integrity in comparison with higher concentration suggesting that at this concentration, antigenic response should be robust.

Since BPL treatment impacted the antigenic potential of S and N, we speculate that it must be causing chemical modifications of amino acids. Similar reports have been made in the case of influenza and coxsackie viruses [6, 30] suggesting that BPL might be interfering with the integrity of the structural proteins of the virion. In agreement with this data, protein quantitation of the BPL treated viral samples always failed to show reproducible results (data not shown). Highly sensitive BCA method detected proteins in the viral samples, albeit in highly irreproducible manner. We demonstrate that viral proteins from such samples could be detected by the Zinc negative staining in SDS-PAGE and could be used for relative quantitation.

Our study demonstrates a clear dichotomy between the Ct values and infectious viral count in any given stock. While the Ct values are representatives of the levels of viral RNA, the plaque assays are direct indicators of the infectivity of the stocks. Our studies demonstrate a lack of correlation between the Ct and PFU values, and a strong correlation between low Ct values and high viral antigens, pointing to the possibility of the presence of a large fraction of non-infectious virus particles in these samples. Having lower infectious titer may not necessarily be a deterrent for samples used in immunizations given that they have high antigenic content. Our results suggest that samples with low Ct and low infectious titers can therefore be used for immunization purposes provided they have good antigenic contents.

While cytopathic effect (CPE) is a clear indication of the viral replication, we observed that several cultures that harbored very high titer virus did not display a CPE during the early part of viral culturing. On the other hand, a few cultures that displayed CPE could not establish viral cultures. Therefore, it is important to monitor the presence of the virus in the supernatant by qRT-PCR. If the cultures do not display CPE, qRT-PCR could be optimally performed at a week from the time of infection.

Our finding on the aggregation of SARS CoV-2 particles has significance in the vaccine and antisera industry. Studies have reported an association between aggregation and loss of antigenicity [31–33]. Another concern is the incomplete inactivation due to aggregation and the potential escape of virus from inactivation resulting in the presence of infectious viruses in vaccine with a potential for infection [34]. However, our studies could not detect any traces of infectious virus after three rounds of consecutive infections. Nevertheless, total inactivation is achieved even at 1:2000 concentrations and hence higher concentrations of BPL are well avoidable.

## CONCLUSION

We have successfully established culturing of multiple isolates of SARS-CoV-2 from patient samples including the modified dry-swab method of collection. We optimized the optimal concentrations of BPL for complete inactivation of SARS-CoV-2. Our studies identified those concentrations of BPL higher than 1:1000 results in aggregation of viral particles and also loss in the antigenic potential of the sample. Our studies would provide a good guiding material for antisera and vaccine studies.

## Author Contributions

D.G. optimized large-scale SARS-CoV-2 virus propagation, BPL inactivation and microneutralization assay. D.G., and D.T propagated, quantified and inactivated large-scale SARS-CoV-2 cultures and performed DLS experiments. D.V. and V.S. established SARS-CoV-2 cultures used in this study. H.P. performed immunoblotting. V.S. performed zinc staining. K.H.H. conceptualized the study and wrote the manuscript.

## Institutional ethics clearance

Institutional ethics clearance (IEC-82/2020) was obtained for the patient sample processing for virus culture.

## Institutional biosafety

Institutional biosafety clearance was obtained for the experiments pertaining to SARS-CoV-2.

## Acknowledgement

We thank several volunteers at the Centre for Cellular and Molecular Biology, who were part of COVID-19 testing that helped us gain access to the potential patient samples for virus culturing. Special thanks to Amit Kumar and Mohan Singh Moodu for their help with the logistics. K Mallesham, R Rukmini helped with DLS experiments and analysis. We thank Karthika Nair, Abhirami P S, Sai Poojitha and Soumya Bunk for their help with experiments.

## Funding

The work was supported by the internal funding from CSIR-CCMB.

## Notes

### Competing Interest Statement

The authors have declared no competing interest.

## References

[1] Lu R, Zhao X, Li J, Niu P, Yang B, Wu H, et al. Genomic characterisation and epidemiology of 2019 novel coronavirus: implications for virus origins and receptor binding. Lancet (London, England). 2020;395:565–74.

[2] Zhu N, Zhang D, Wang W, Li X, Yang B, Song J, et al. A Novel Coronavirus from Patients with Pneumonia in China, 2019. The New England journal of medicine. 2020;382:727–33.

[3] Puelles VG, Lütgehetmann M, Lindenmeyer MT, Sperhake JP, Wong MN, Allweiss L, et al. Multiorgan and Renal Tropism of SARS-CoV-2. The New England journal of medicine. 2020;383:590–2.

[4] Chen H, Wu R, Xing Y, Du Q, Xue Z, Xi Y, et al. Influence of Different Inactivation Methods on Severe Acute Respiratory Syndrome Coronavirus 2 RNA Copy Number. Journal of clinical microbiology. 2020;58.

[5] Furuya Y, Chan J, Regner M, Lobigs M, Koskinen A, Kok T, et al. Cytotoxic T cells are the predominant players providing cross-protective immunity induced by {gamma}-irradiated influenza A viruses. Journal of virology. 2010;84:4212–21.

[6] Fan C, Ye X, Ku Z, Kong L, Liu Q, Xu C, et al. Beta-Propiolactone Inactivation of Coxsackievirus A16 Induces Structural Alteration and Surface Modification of Viral Capsids. Journal of virology. 2017;91.

[7] Roberts A, Lamirande EW, Vogel L, Baras B, Goossens G, Knott I, et al. Immunogenicity and protective efficacy in mice and hamsters of a β-propiolactone inactivated whole virus SARS-CoV vaccine. Viral immunology. 2010;23:509–19.

[8] Patterson EI, Prince T, Anderson ER, Casas-Sanchez A, Smith SL, Cansado-Utrilla C, et al. Methods of Inactivation of SARS-CoV-2 for Downstream Biological Assays. The Journal of infectious diseases. 2020;222:1462–7.

[9] Jureka AS, Silvas JA, Basler CF. Propagation, Inactivation, and Safety Testing of SARS-CoV-2. Viruses. 2020;12.

[10] Kariwa H, Fujii N, Takashima I. Inactivation of SARS coronavirus by means of povidone-iodine, physical conditions, and chemical reagents. Jpn J Vet Res. 2004;52:105–12.

[11] See RH, Petric M, Lawrence DJ, Mok CPY, Rowe T, Zitzow LA, et al. Severe acute respiratory syndrome vaccine efficacy in ferrets: whole killed virus and adenovirus-vectored vaccines. The Journal of general virology. 2008;89:2136–46.

[12] See RH, Zakhartchouk AN, Petric M, Lawrence DJ, Mok CPY, Hogan RJ, et al. Comparative evaluation of two severe acute respiratory syndrome (SARS) vaccine candidates in mice challenged with SARS coronavirus. The Journal of general virology. 2006;87:641–50.

[13] Kempner ES. Effects of high-energy electrons and gamma rays directly on protein molecules. Journal of pharmaceutical sciences. 2001;90:1637–46.

[14] Kiran U, Gokulan CG, Kuncha SK, Vedagiri D, Chander BT, Sekhar AV, et al. Easing diagnosis and pushing the detection limits of SARS-CoV-2. Biology methods & protocols. 2020;5:bpaa017.

[15] George A, Panda S, Kudmulwar D, Chhatbar SP, Nayak SC, Krishnan HH. Hepatitis C virus NS5A binds to the mRNA cap-binding eukaryotic translation initiation 4F (eIF4F) complex and up-regulates host translation initiation machinery through eIF4F-binding protein 1 inactivation. The Journal of biological chemistry. 2012;287:5042–58.

[16] Schneider CA, Rasband WS, Eliceiri KW. NIH Image to ImageJ: 25 years of image analysis. Nature methods. 2012;9:671–5.

[17] Kaye M. SARS-associated coronavirus replication in cell lines. Emerging infectious diseases. 2006;12:128–33.

[18] Stelzer-Braid S, Walker GJ, Aggarwal A, Isaacs SR, Yeang M, Naing Z, et al. Virus isolation of severe acute respiratory syndrome coronavirus 2 (SARS-CoV-2) for diagnostic and research purposes. Pathology. 2020;52:760–3.

[19] Caly L, Druce J, Roberts J, Bond K, Tran T, Kostecki R, et al. Isolation and rapid sharing of the 2019 novel coronavirus (SARS-CoV-2) from the first patient diagnosed with COVID-19 in Australia. The Medical journal of Australia. 2020;212:459–62.

[20] Díaz FJ, Aguilar-Jiménez W, Flórez-Álvarez L, Valencia G, Laiton-Donato K, Franco-Muñoz C, et al. Isolation and characterization of an early SARS-CoV-2 isolate from the 2020 epidemic in Medellín, Colombia. Biomedica: revista del Instituto Nacional de Salud. 2020;40:148–58.

[21] Pujadas E, Chaudhry F, McBride R, Richter F, Zhao S, Wajnberg A, et al. SARS-CoV-2 viral load predicts COVID-19 mortality. The Lancet Respiratory medicine. 2020;8:e70.

[22] Karahasan Yagci A, Sarinoglu RC, Bilgin H, Yanılmaz Ö, Sayin E, Deniz G, et al. Relationship of the cycle threshold values of SARS-CoV-2 polymerase chain reaction and total severity score of computerized tomography in patients with COVID 19. International journal of infectious diseases: IJID: official publication of the International Society for Infectious Diseases. 2020;101:160–6.

[23] Borsanyiova M, Kubascikova L, Sarmirova S, Vari SG, Bopegamage S. Assessment of a swab collection method without virus transport medium for PCR diagnosis of coxsackievirus infections. Journal of virological methods. 2018;254:18–20.

[24] Moore C, Corden S, Sinha J, Jones R. Dry cotton or flocked respiratory swabs as a simple collection technique for the molecular detection of respiratory viruses using real-time NASBA. Journal of virological methods. 2008;153:84–9.

[25] La Scola B, Le Bideau M, Andreani J, Hoang VT, Grimaldier C, Colson P, et al. Viral RNA load as determined by cell culture as a management tool for discharge of SARS-CoV-2 patients from infectious disease wards. European journal of clinical microbiology & infectious diseases: official publication of the European Society of Clinical Microbiology. 2020;39:1059–61.

[26] Kim SE, Jeong HS, Yu Y, Shin SU, Kim S, Oh TH, et al. Viral kinetics of SARS-CoV-2 in asymptomatic carriers and presymptomatic patients. International journal of infectious diseases: IJID: official publication of the International Society for Infectious Diseases. 2020;95:441–3.

[27] Wiktor TJ, Aaslestad HG, Kaplan MM. Immunogenicity of rabies virus inactivated by -propiolactone, acetylethyleneimine, and ionizing irradiation. Applied microbiology. 1972;23:914–8.

[28] Logrippo GA, Hartman FW. Antigenicity of beta-propiolactone-inactivated virus vaccines. Journal of immunology (Baltimore, Md: 1950). 1955;75:123–8.

[29] Perrin P, Morgeaux S. Inactivation of DNA by beta-propiolactone. Biologicals: journal of the International Association of Biological Standardization. 1995;23:207–11.

[30] She YM, Cheng K, Farnsworth A, Li X, Cyr TD. Surface modifications of influenza proteins upon virus inactivation by β-propiolactone. Proteomics. 2013;13:3537–47.

[31] Kon TC, Onu A, Berbecila L, Lupulescu E, Ghiorgisor A, Kersten GF, et al. Influenza Vaccine Manufacturing: Effect of Inactivation, Splitting and Site of Manufacturing. Comparison of Influenza Vaccine Production Processes. PloS one. 2016;11:e0150700.

[32] Desbat B, Lancelot E, Krell T, Nicolaï M-C, Vogel F, Chevalier M, et al. Effect of the β-propiolactone Treatment on the Adsorption and Fusion of Influenza A/Brisbane/59/2007 and A/New Caledonia/20/1999 Virus H1N1 on a Dimyristoylphosphatidylcholine/Ganglioside GM3 Mixed Phospholipids Monolayer at the Air–Water Interface. Langmuir. 2011;27:13675–83.

[33] Chen Y, Zhang Y, Quan C, Luo J, Yang Y, Yu M, et al. Aggregation and antigenicity of virus like particle in salt solution--A case study with hepatitis B surface antigen. Vaccine. 2015;33:4300–6.

[34] Delrue I, Verzele D, Madder A, Nauwynck HJ. Inactivated virus vaccines from chemistry to prophylaxis: merits, risks and challenges. Expert review of vaccines. 2012;11:695–719.

